# Accuracy of demographic inferences from site frequency spectrum: the case of the yoruba population

**DOI:** 10.1101/078618

**Authors:** Marguerite Lapierre, Amaury Lambert, Guillaume Achaz

## Abstract

Some methods for demographic inference based on the observed genetic diversity of current populations rely on the use of summary statistics such as the Site Frequency Spectrum (SFS). Demographic models can be either model-constrained with numerous parameters such as growth rates, timing of demographic events and migration rates, or model-flexible, with an unbounded collection of piecewise constant sizes. It is still debated whether demographic histories can be accurately inferred based on the SFS. Here we illustrate this theoretical issue on an example of demographic inference for an African population. The SFS of the Yoruba population (data from the 1000 Genomes Project) is fit to a simple model of population growth described with a single parameter (*e.g.*, founding time). We infer a time to the most recent common ancestor of 1.7 million years for this population. However, we show that the Yoruba SFS is not informative enough to discriminate between several different models of growth. We also show that for such simple demographies, the fit of one-parameter models outperforms the model-flexible method recently developed by Liu and Fu. The use of this method on simulated data suggests that it is biased by the noise intrinsically present in the data.

## INTRODUCTION

Inference of human population history based on demographic models for genomic data can complement archaeological knowledge, owing to the large amount of polymorphism data now available in human populations. Polymorphism data can be viewed as an imprint left by past demographic events on the current genetic diversity of a population (see, *e.g.*, review by POOL *et al*. 2010).

There are several means of analyzing this observed genetic diversity for demographic inference. The polymorphism data can be used to reconstruct a coalescence tree of the sampled individuals. The demography of the sampled population can be inferred by comparing this reconstructed tree with theoretical predictions under a constant size model (PYBUS *et al*. 2000). For example, in an expanding population, the reconstructed coalescent tree will have relatively longer terminal branches than the reference coalescent tree in a population of constant size. However, methods based on a single reconstructed tree are flawed because of recombination (LAPIERRE *et al*. 2016), since the genealogy of a recombining genome is described by as many trees as there are recombining loci.

The genome-wide distribution of allele frequencies is a function of the average genealogies, and can thus be used as a summary statistic for demographic inference. This distribution, called the Site Frequency Spectrum (SFS), reports the number of mutated sites at any given frequency. The demographic history of a population affects the shape of its SFS (ADAMS and HUDSON 2004; MARTH *et al*. 2004). For example, an expanding population carries an excess of low-frequency variants, compared with the expectation under a constant size model. The shape of the SFS is also altered by selection, which results in an excess of low- and high-frequency variants (FAY and WU 2000). However, selection acts mainly on the coding parts of the genome and the non-coding segments linked to them, while demography impacts the whole genome. Furthermore, unlike reconstructed trees, the SFS is not biased by recombination (WALL 1999). Quite on the contrary, by averaging the SFS over many correlated marginal genealogies, recombination lowers the variance of the SFS while its expectation remains unchanged. Therefore, the SFS of a sample is a summary of the genetic diversity, averaged over all the genome due to recombination, that can be analyzed in terms of demography.

Several types of methods exist to infer the demography of a population based on its SFS. A specific demographic model can be tested by computing a pseudo-likelihood function for this model, based on the comparison of the observed SFS and the SFS estimated by Monte Carlo coalescent tree simulations (NIELSEN 2000; COVENTRY *et al*. 2010; NELSON *et al*. 2012). This method can be extended to infer demographic scenarios of several populations, using their joint SFS (EXCOFFIER *et al*. 2013). Methods based on Monte Carlo tree simulations are typically very costly in computation time. Other approaches rely on diffusion processes: they use the solution to the partial differential equation of the density of segregating sites as a function of time (GUTENKUNST *et al*. 2009; LUKIć *et al*. 2011).

Whereas all these methods are model-constrained, *i.e.*, they use the SFS to test the likelihood of a given demographic model, more flexible methods have been developed. Recently, BHASKAR *et al*. (2015) derived exact expressions of the expected SFS for piecewise-constant and piecewise-exponential demographic models. LIU and FU (2015) developed a model-exible method based on the SFS: the stairway plot. This method infers the piecewise-constant demography which maximizes the composite likelihood of the SFS, without any previous knowledge on the demography. This optimization is based on the estimation of a time-dependent population mutation rate *θ*. Although they show that their method infers efficiently some theoretical demographies, they do not test the goodness of fit of the expected SFS, reconstructed under the demography they infer, with the input SFS on which they apply their method.

All these methods are widely used for the inference of demography in humans and other species, but doubts remain on the identifiability of a population demography based on its SFS. It has been shown theoretically that certain population size functions are unidentifiable from the population SFS (MYERS *et al*. 2008; TERHORST and SONG 2015). MYERS *et al*. (2008) showed that for any given population size function *N*(*t*), there exists an infinite number of smooth functions *F*(*t*) such that *ξ*^N^ = *ξ*^N+F^ where *ξ*^N^ is the SFS of a population of size function *N*(*t*). However, other theoretical works have recently shown that for many types of population size functions commonly used in demography studies, such as piecewise constant or piecewise exponential functions, demography can be inferred based on the SFS, provided the sample is large enough (BHASKAR and SONG 2014). These studies argued that the unidentifiability proven by MYERS *et al*. (2008) relied on biologically unrealistic population size functions involving high frequency oscillations near the present. Lately, two studies (KIM *et al*. 2015; TERHORST and SONG 2015) have provided bounds on the amount of demographic information contained in the SFS or in coalescent times.

In this study, we use the SFS of an African population (the Yoruba population, data from The 1000 Genomes Project Consortium 2015) as an example of a somewhat simple demography, to illustrate the risks of over-confidence in demographic scenarios inferred. Namely, we highlight two issues potentially arising even in the case of simple demographies: unidentifiability of models and poor goodness of fit of inferences. We first infer the Yoruba demography with a model-constrained method, using diverse one-parameter models of growth, and then with a model-flexible method, the stairway plot (Liu and Fu 2015). For the model-constrained method, we test four different growth models derived from the standard neutral framework used in the vast majority of population genetics studies, also compared with a more uncommon type of model based on a branching process. Individual-based models such as the branching process are widely used in population ecology (LAMBERT 2010): the population is modeled as individuals which die and give birth at given rates independently. These models are not commonly used in population genetics although they provide interesting features of fluctuating population sizes for example, and benefit from a strong mathematical framework.

## MATERIALS AND METHODS

**1000 Genomes Project data:** Variant calls from the 1000 Genomes Project phase 3 were downloaded from the project ftp site (THE 1000 GENOMES PROJECT CONSORTIUM 2015). The sample size for the Yoruba population is *n* = 108 individuals (polymorphism data available for both genome copies of each individual, *i.e.*, 2*n* = 216 sequences). We kept all single nucleotide bi-allelic variants to plot the sample SFS. The number of bi-allelic sites is *S* = 20417698. The average distance between two sites is 136 bp (median 81 bp). The number of sites for which the ancestral allele is known is *S*^'^ = 19441528. To avoid possible bias due to sequencing errors, we ignored singletons (mutations appearing in only one chromosome of one individual in the sample) for the rest of the study. The implications of ignoring singletons are examined in the discussion.

**Site Frequency Spectrum definition and graphical representation:** The Site Frequency Spectrum (SFS) of a sample of *n* diploid individuals is described as the vector *ξ* = (*ξ*_1_, *ξ*_2_, …, *ξ*_2*n*–1_) where for *i* ∈ [1, 2*n*−1], *ξ*_*i*_ is the number of dimorphic (*i.e.*, with exactly two alleles) sites with derived form at frequency *i*/2*n*. To avoid potential orientation errors, we assumed that the ancestral form is unknown for all sites: we worked with a folded spectrum, where we consider the frequency of the less frequent (or minor) allele. In this case, the folded SFS is described as the vector *η* = (*η*_1_, *η*_2_,…, *η*_*n*_) where η_*i*_ = *ξ*_*i*_ + *ξ*_2*n* − *i*_ for *i* ∈ [1, *n* − 1] and η_*n*_ = *ξ*_*n*_. The folded SFS of the Yoruba sample is plotted in Figure S1. For a better graphical representation, all SFS were transformed as follows: we plot *ϕ*_*i*_ normalized by its sum, where

- for unfolded SFS,

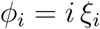 for *i* ∈ [1, 2*n*-1]
- for folded SFS,

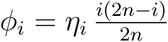 for *i* ∈ [1, *n*-1] and *ϕ*_*n*_ = *n* η_*n*_

The transformed SFS has a flat expectation (*i.e.*, constant over all values of *i*) under the standard neutral model (NAWA and TAJIMA 2008; ACHAZ 2009).

**Demographic models used for the model-constrained methods**: We inferred the demography of the Yoruba population using five growth models (Figure 1), compared with the predictions of the standard model with constant population size. Time is measured in coalescent units of 2*N* generations, where the scaling parameter *N* has the same dimension as the current population size, which we will not estimate. Time starts at 0 (present time) and increases backward in time. Four models are based on the standard Kingman coalescent (KINGMAN 1982), amended with demography. Three of them are described with an explicit demography: either *Linear* growth since time *τ*, *Exponential* growth at rate 1/*τ* or *Sudden* growth from a single ancestor to the entire population at time *τ*. We also use another model based on the Kingman coalescent, with an implicit demography: the *Conditioned* model. This model is based on a standard constant size model, but the Time to the Most Recent Common Ancestor (*T*_*MRCA*_) is conditioned on being reached before time *τ*. The fifth model, *Birth-Death*, is not based on the standard Kingman coalescent, but on a critical branching process measured in units of 2*N* generations. Forward in time, the process starts with a founding event of one individual. Individuals give birth and die at equal rate 1. The process is conditioned on not becoming extinct before a period of time *τ*, and on reaching on average 2*N* individuals.

**Figure 1:**
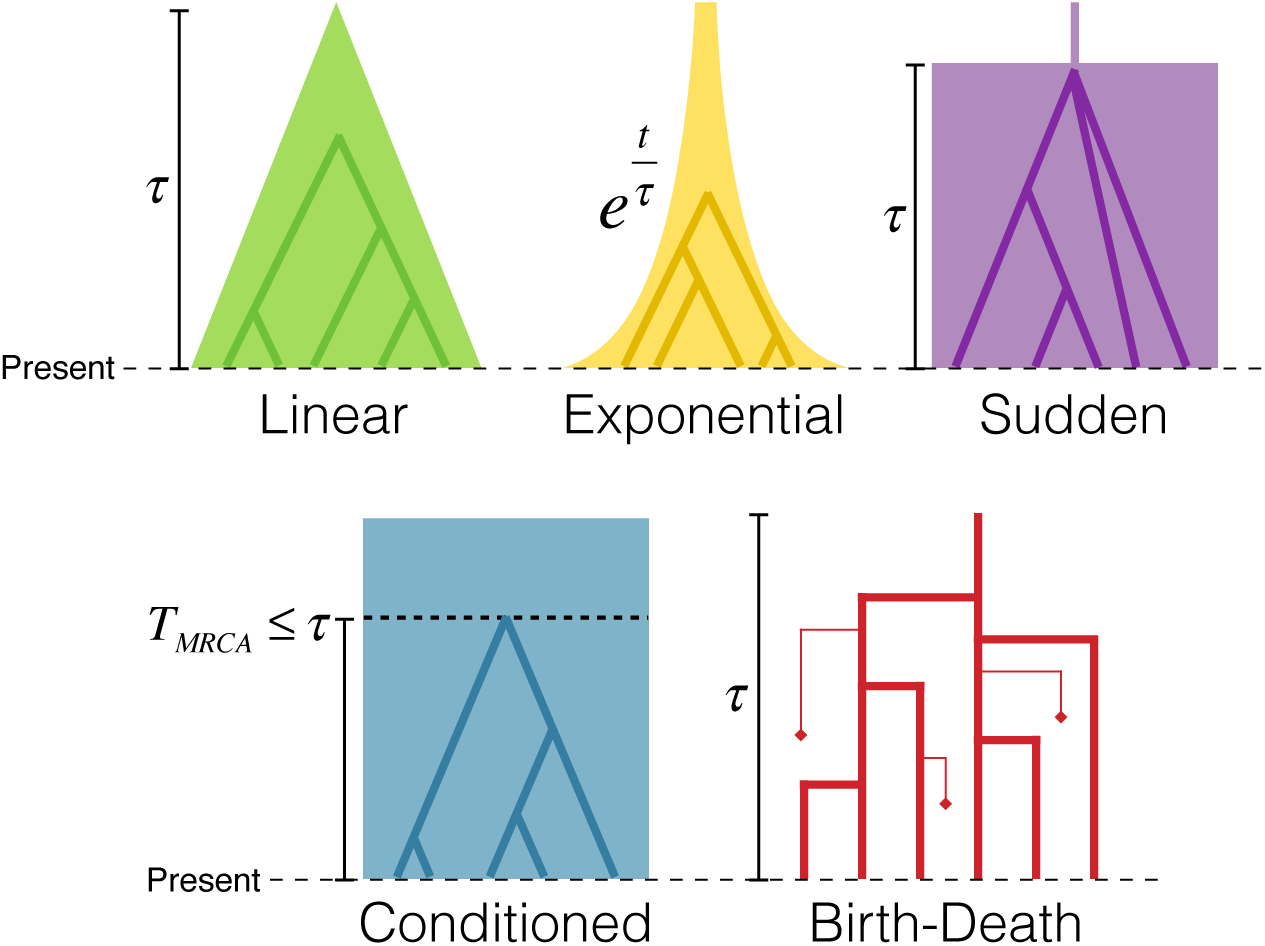
The five demographic models. Each model has one single time parameter *τ*.

**Stairway plot inference on the Yoruba SFS:** We applied the model-flexible stairway plot method developed by LIU and FU (2015) on the unfolded Yoruba SFS. Inferences are made on 200 SFS as suggested by their method. We use the script they provide to create 199 bootstrap samples of the Yoruba SFS. We also ignore the singletons for this method, and use the default parameter values suggested in their paper for the optimization.

**SFS simulation with demography:** We used two different method to simulate SFS under the four demographic models derived from the Kingman coalescent (*Linear, *Exponential*, Sudden* and *Conditioned*) or under a piecewise-constant demography reconstructed by the stairway plot method.

*Method 1:* Simulate *l* independent topologies under the Kingman coalescent on which mutations are placed at rate *θ* (population mutation rate) (HUDSON *et al*. 1990). This allows to simulate the SFS of *l* independent loci.

*Method 2:* Another way to simulate SFS is using the following formula:

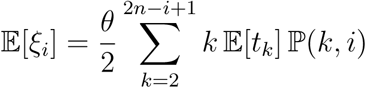

where *θ* is the population mutation rate, *t*_*k*_ is the time during which there are *k* lines in the tree (hereafter named state *k*) and ℙ (*k*, *i*) is the probability that a randomly chosen line at state *k* gives *i* descendants in the sample of size 2*n* (*i.e.*, at state 2*n*) (FU 1995). For all models, the neutrality assumption ensures that

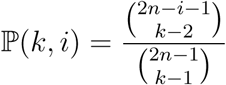

for *i* ∈ [1, 2*n* − 1] and *k* ∈ [2, 2*n*−*i* + 1]. Using this probability allows to average over the space of topologies. This reduces considerably computation time since the space of topologies is very large, and produces smooth SFS for which only the *t*_*k\**_ need to be simulated to obtain the expectations
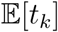.

The expectations 
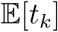
 are obtained as follow: for *k* ∈ [2, 2*n*], times in the standard coalescent *t*_*k*_ are drawn from an exponential distribution of parameter
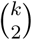
. For the *Linear* and *Exponential* models, and for the piecewise-constant demographies reconstructed by the stairway plot method, these times are then rescaled to take into account the given explicit demography (see, *e.g.*, HEIN *et al*. 2004, chap.4). For the *Sudden* model, we assume the coalescence of all lineages at time *τ* if the common ancestor has not been reached yet. For the *Conditioned* model, we keep only simulations for which 
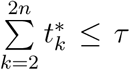
 where *τ* is the model parameter. The expectations 
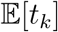
 are obtained by averaging over 10^7^ simulations.

Alternatively, the expectations 
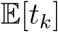 could also be obtained with analytic formulae provided by POLANSKI and KIMMEL (2003).

For the *Birth-Death* model, we use the explicit formula for the SFS given in DELAPORTE *et al*. (2016).

We normalize the SFS computed under all these models so that their sum equals 1. This normalization removes the dependence on the mutation rate parameter *θ*. Consequently, the standard model has no parameters while all others have exactly one (*τ*).

**Optimization of the parameter *τ*:** For each demographic model, we optimize the parameter by minimizing the weighted square distance *d*^2^ between the observed SFS of the Yoruba population and the predicted SFS under the model (simulated with *Method 2*). Both SFS are normalized for comparison. The distance is computed for all *τ* values in a given interval (no specific optimization method was used to find the minimum). With 
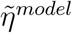
 and 
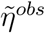
 the folded and normalized SFS in the tested model and in the data respectively,

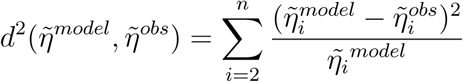

The sum starts at *i* = 2 because we ignore 
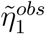
 corresponding to singletons. To calculate the distance *d*^2′^ between the SFS predicted by two models A and B, we weight the terms by the mean of the two models:

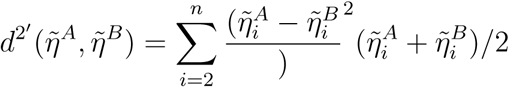

**nference of the Yoruba demography with** ∂**a**∂**i:** We inferred the demography of the Yoruba population with the software ∂a∂i v1.7 (GUTENKUNST *et al*. 2009), testing the three models of explicit demography (*Linear*, *Exponential* and *Sudden*). The demographic models were specified so that the only parameter to optimize is *τ* like for the distance-based inference method. Singletons were masked and the method was applied on the folded Yoruba SFS. Details on the demographic functions and parameter values used for the optimization in ∂a∂i are provided in the Supplementary Methods. We ran the method 100 times for each model and kept the parameter value with the best maximum log composite likelihood over the 100 runs. In Figure S4, we plot the best log composite likelihood of the 100 runs.

**Scaling of the coalescent time:** Optimized values of the parameter 
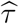
 for each model are expressed in coalescent time units, *i.e.*, scaled in 2*N*(0) generations. As the model population size at time zero, 2*N*(0), is unknown, to scale these coalescent time units in numbers of generations and consequently in years, we used the expected number of mutations per site *M*. From the dataset, we have *M*^*obs*^ = *S*/*L* where *S* is the number of single nucleotide mutations (a *k*-allelic SNP accounts for *k* − 1 mutations) and *L* is the length of the accessible sequenced genome in the 1000 genomes project (90% of the total genome length, THE 1000 GENOMES PROJECT CONSORTIUM). For the theoretical value, we get that 
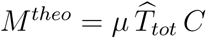
 where we know the mutation rate *μ* from the literature and the total tree length expressed in coalescent time units 
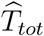
 from the SFS simulations. Here *C* is the coalescent factor, that is the number of generations per coalescent time unit, also corresponding to 2*N*_*e*_(0) where *N*_*e*_(0) is the effective population size at present time. The total number of generations in the tree is 
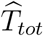
 *C* from which we derive the total number of mutations per site *M*^*theo*^. Thus, using the observed value *M*^*obs*^, we can estimate *C* by 
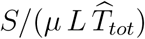
 We assumed a mutation rate of 1.2 × 10 ^−8^ per base pair per generation (CONRAD *et al*. 2011; CAMPBELL *et al*. 2012; KONG *et al*. 2012). With the coalescent factor *C*, we can then convert a coalescent time unit into a number of generations, or into a number of years assuming 24 years as generation time (SCALLY and DURBIN 2012).

**Graphical representation of the inferred demographies**: To represent the inferred explicit demographies (models *Linear, Exponential* and *Sudden*), we plot the shape of the demography with the optimized value 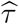
 for each model. For the implicit demographies (models *Conditioned* and *Birth-Death*), as there is no explicit demographic shape, we plot the mean trajectory of fixation of a new allele in the population: forward in time, these fixation trajectories illustrate the expansion of the descendance of the sample’s ancestor in the population (see the Supplementary Methods for details).

**Comparing the model-constrained and model-flexible methods to infer *Linear* growth:** We applied both methods (the one-parameter inference method and the stairway plot method) on SFS simulated under *Linear* growth. To test the stairway plot method on a *Linear* growth demography, we simulate 200 independent SFS using *Method 1*, with sample size 2*n* = 216, *θ* = 100 (arbitrary value removed by normalization) and a founding time *τ* = 2.48 (estimated for the Yoruba population, see Results). The SFS are simulated with either 10^3^, 10^4^ or 10^5^ independent loci. We scaled the simulated SFS to obtain a total number of *S* = 20417698 variants, so that the total number of variants in the simulated SFS is the same as in the Yoruba SFS. We ran the stairway plot method on these 200 independent SFS with the default parameter values suggested in the method, and with the same mutation rate (1.2 × 10 ^−8^ per base pair per generation) and generation time (24 years) as in our study. We report the median demography of these 200 independent inferences.

To test the one-parameter inference method on these SFS simulated under the *Linear* model, we run the parameter optimization on a SFS simulated with either 10^3^, 10^4^, 10^5^ or 10^6^ loci. The search of the parameter value that minimizes the distance *d*^2^ was optimized with a Newton-Raphson algorithm. Derivatives were calculated at *τ* ± 0.05 where *τ* is the parameter value being optimized. The optimization stopped when the optimization step of the parameter value was smaller than 10 ^−3^.

**Data and software availability** The 1000 genomes project data used in this study is publicly available at ftp://ftp.1000genomes.ebi.ac.uk/vol1/ftp/release/20130502/. The code in Python and C written for the study is available athttps://github.com/lapierreM/Yoruba_demography. The code in C used for the *Method 1* of SFS simulation is available upon request to G. ACHAZ.

## RESULTS

We inferred the demography of the Yoruba population (Africa), from the whole-genome polymorphism data of 108 individuals (data from the 1000 Genomes Project, THE 1000 GENOMES PROJECT CONSORTIUM), with SFS-based methods, either model-constrained or model-flexible.

It has been shown that human populations have been growing since their emergence in Africa, and that African populations were supposedly not affected by the Out-of-Africa bottleneck described for Eurasian populations (MARTH *et al*. 2004; ATKINSON *et al*. 2008; GUTENKUNST *et al*. 2009; GRONAU *et al*. 2011; TENNESSEN *et al*. 2012). Analyses using the PSMC method (LI and DURBIN 2011) have shown a reduction of the African population size after the divergence with non-African populations. However, MAZET *et al*. (2016) have recently shown that these analyses could be biased by population structure. Based on this previous knowledge, for the model-constrained method, we chose to infer the Yoruba demography with simple models of growth, *i.e.*, with only one phase of growth characterized by a single parameter. These five models are: *Linear, Exponential* or *Sudden* growth, a *Conditioned* model where the *T*_*MRCA*_ is conditioned on being smaller than the given parameter, and a critical *Birth-Death* model based on a branching process (Figure 1). To infer the Yoruba demography, we fit the SFS predicted under each model with the observed Yoruba SFS (all SFS are folded). The SFS were normalized to remove the population mutation rate parameter *θ*, so that each model is characterized by one single parameter *τ* which has the dimension of a time duration. We fit this parameter by least-square distance between the observed SFS and the predicted SFS, and by maximum likelihood using the ∂a∂i software (GUTENKUNST *et al*. 2009). For the model-flexible inference, we used the stairway plot method developed recently by LIU and FU (2015), which infers a piecewise-constant demography based on the SFS. For this method, the number of parameters to be estimated is determined by a likelihood-ratio test. It can range from 1 to 2*n* − 1 where 2*n* is the number of sequences in the sample.

The Yoruba SFS was constructed by taking into account the entire genome. Removing the coding parts of the genome to avoid potential bias due to selection does not affect the shape of the SFS substantially (Figure S2), since the coding parts represent a very small fraction of the human genome. The first bin of the observed SFS, accounting for mutations found in one chromosome of one individual in the sample (black dot in the observed SFS in Figure 3B), seemed to lie outside the rest of the distribution. This could be due to sequencing errors being considered as singletons (ACHAZ 2008), and thus we chose to ignore this value for the model optimization. We have also made sure that the SFS shape was not affected greatly by the sample size. We compared the SFS of a subsample of half the Yoruba individuals (2*n* = 108) with the full sample SFS (2*n* = 216) (Figure S3). This shows that the only bin of the SFS which is significantly affected by this subsampling is the first one, containing the singletons. As we ignore it in our study, it does not influence our results.

The analysis of the Yoruba SFS with the stairway plot method results in a complex demography with several bottlenecks in the last 160 000 years (Figure 2). The current effective population size *N_e_*(0) is 28 500 (time 0 does not correspond to present time as we ignored singletons, see discussion). The demographic history earlier than 160 000 years ago shows spurious patterns that should not be interpreted, according to LIU and FU (2015).

**Figure 2:**
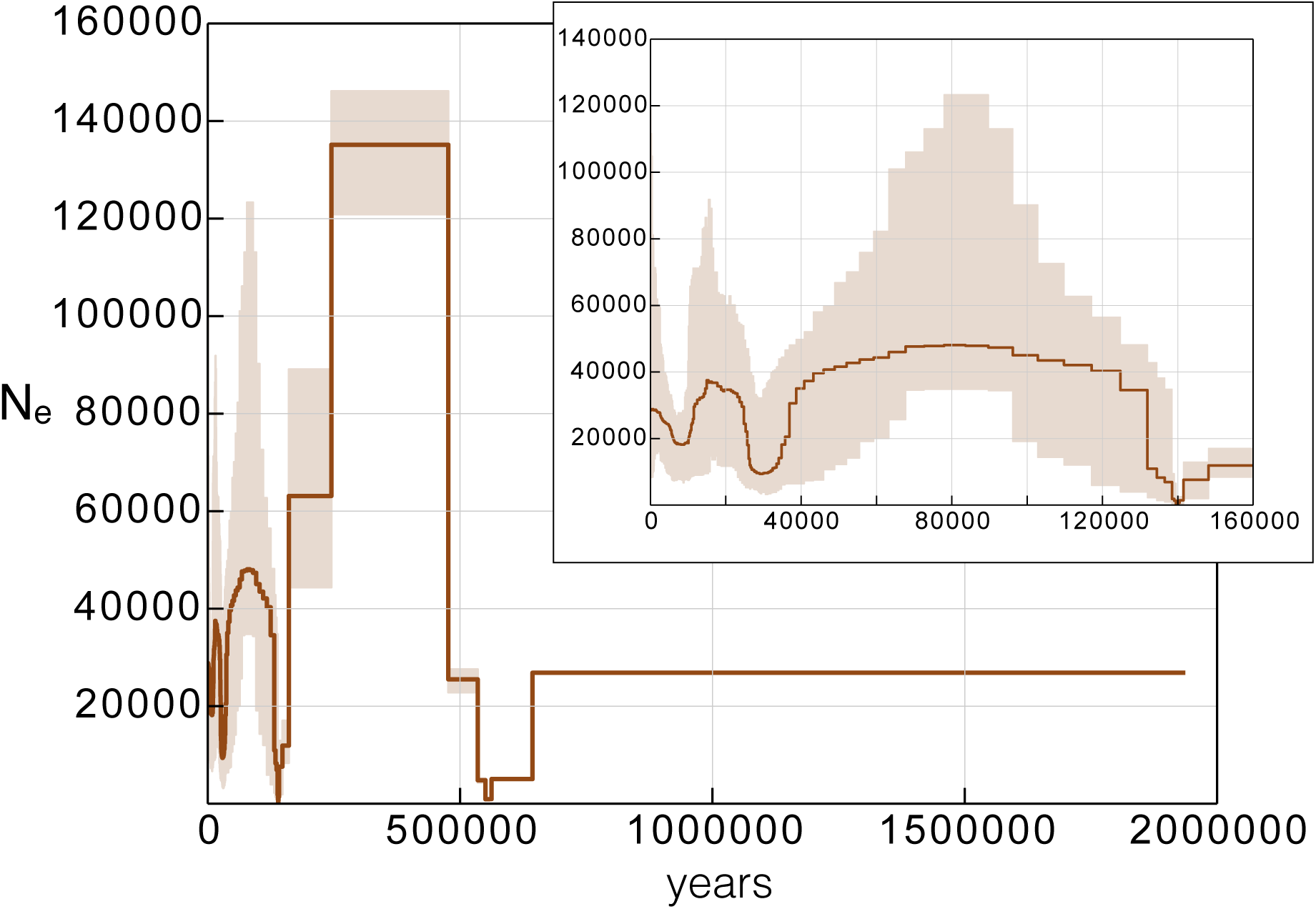
Stairway plot inference of the Yoruba demography. The inferred effective size *N_e_* of the Yoruba population is plotted from present time (0) to the past. The inset is a zoom between 0 and 160 000 years. The thick brown line is the median *N_e_*, the light brown area is the [2.5; 97.5] percentiles interval. The inference is based on 200 bootstrap samples of the unfolded Yoruba SFS. The singletons are not taken into account for the optimization of the stairway plot.

The inference of the Yoruba demography with one-parameter models was done by minimizing the distance between observed and predicted SFS. This gave an optimized value 
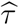
 of the parameter τ (Figure 3A and Table 1) (with 
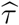
 in coalescent units, *Linear*: 
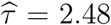
 *Exponential*: 
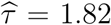
 *Sudden*: 
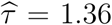
 *Conditioned*: 
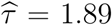
 *Birth-Death*: 
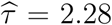
 Plotting the predicted SFS with the optimized parameter value 
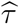
 confirmed their goodness of fit with the observed Yoruba SFS (Figure 3B). Compared to the standard model without demography, the addition of just one parameter allows for a surprisingly good fit of the observed Yoruba SFS. The Yoruba demography thus seems to be compatible with a simple scenario of growth. On the other hand, the demography inferred by the stairway plot predicts a SFS which does not fit well the observed Yoruba SFS: the distance between the observed Yoruba SFS and the expected SFS under the stairway plot demography is ten times the distance between any of the one-parameter model SFS and the data (Figure 3B and Table 1).

**Figure 3:**
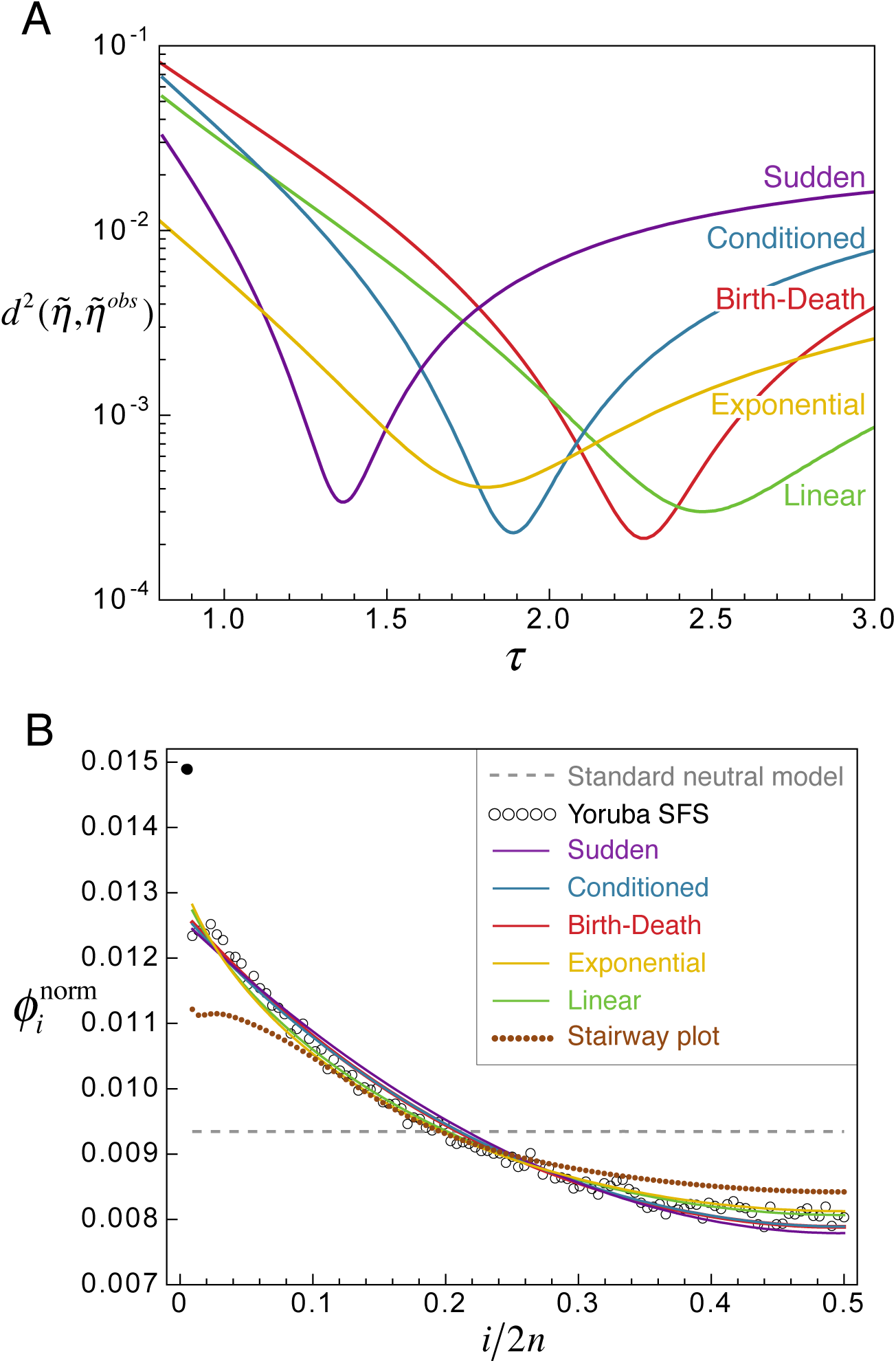
Inference of the Yoruba demography with one-parameter models. A) Weighted square distance 
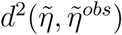
 between the normalized Yoruba SFS 
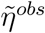
 and the normalized predicted SFS under each of the five models, depending on the value of the parameter *τ* (Purple: *Sudden*, Blue: *Conditioned*, Red: *Birth-Death*, Yellow: Exponential, Green: *Linear*). B) Predicted SFS under each of the five models, with the optimized value 
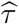
 of the parameter, and under the demography inferred by the stairway plot (brown dotted line). The Yoruba SFS is shown in empty circles. The first dot, colored in black, accounting for the singletons, was not taken into account for the optimization of *τ* to avoid potential bias due to sequencing errors. The grey dashed line is the expected SFS under the standard neutral model without demography. Colors match the plot above (the predicted SFS under the models *Birth-Death* and *Conditioned* are indistinguishable). The SFS are folded, transformed and normalized (see Methods).

**Table 1:** Least-square distance *d*^2^ between pairs of observed Yoruba SFS and optimized SFS under the five demographic models or the stairway plot method.

|  | Data | *Linear* | *Exponential* | *Sudden* | *Conditioned* | *Birth-Death* |
| --- | --- | --- | --- | --- | --- | --- |
| *Linear* | $3.0 \times 10^{-4}$ | 0 | | | | |
| *Exponential* | $4.1 \times 10^{-4}$ | $2.2 \times 10^{-5}$ | 0 | | | |
| *Sudden* | $3.4 \times 10^{-4}$ | $3.5 \times 10^{-4}$ | $5.5 \times 10^{-4}$ | 0 | | |
| *Conditioned* | $2.3 \times 10^{-4}$ | $1.6 \times 10^{-4}$ | $5.5 \times 10^{-4}$ | $3.7 \times 10^{-5}$ | 0 | |
| *Birth-Death* | $2.2 \times 10^{-4}$ | $1.7 \times 10^{-4}$ | $3.1 \times 10^{-4}$ | $4.1 \times 10^{-5}$ | $3.5 \times 10^{-6}$ | 0 |
| Stairway plot | $2.9 \times 10^{-3}$ | $3.1 \times 10^{-3}$ | $3.3 \times 10^{-3}$ | $2.8 \times 10^{-3}$ | $2.8 \times 10^{-3}$ | $2.9 \times 10^{-3}$ |

The best fitting SFS under each of the five demographic models all have a square distance *d*^2^ of the order of 10 ^−4^ with the observed Yoruba SFS (Figure 3A and Table 1) and have highly similar shapes (Figure 3B). This suggests that the ve demographic models used to infer the demography of the Yoruba are hard to distinguish based only on the observed SFS. To validate the use of a least square distance to find the best fitting SFS, we also infered the Yoruba demography using the ∂a∂i software. This model-constrained method based on the SFS uses a diffusion approximation to simulate SFS and a likelihood framework for the parameter optimization. We tested the three models of explicit demography (*Linear*, *Exponential* and *Sudden* growth) parametrized in the same way as in our method. The best parameter values found by ∂a∂i by maximum log composite likelihood are the same as by our method (with 
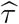
 in coalescent units, *Linear*: 
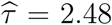
 *Exponential*: 
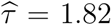
 *Sudden*: 
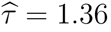
. Moreover, the log composite likelihoods of the best fitting SFS under each model are on the same scale (the likelihoods are directly comparable because the number of parameters is the same for each model): *Linear*: ln(*L*) = −3107, *Exponential*: ln(*L*) = −3953, *Sudden*: ln(*L*) =−3393 (Figure S4). They rank the explicit demography models in the same order as the least square distance *d*^2^ would rank them: the best model is *Linear* growth, then *Sudden* and finally *Exponential* growth.

We computed the expected *T*_*MRCA*_ based on the predicted SFS using (1): as the SFS predicted under each model are very similar, it means that they have roughly the same estimated time durations *t*_*k*_ while there are *k* branches in the coalescent tree of the Yoruba sample. From these expected *t*_*k*_ we can compute 
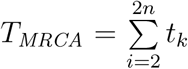
 This is the *T*_*MRCA*_ of the sample, but we can assume that it is the same as the *T*_*MRCA*_ of the population, because with such a large sample size, the probability that the *T*_*MRCA*_ of the population is different from the *T_MRCA_* of the sample becomes very small. Under each of four models (excluding the *Birth-Death* model for which there is no obvious common time scaling), the expected *T*_*MRCA*_ for the Yoruba population is 1.3 in coalescent units. By using the number of mutations per site in the data and the total tree length inferred from the simulations, we scaled back this *T*_*MRCA*_ in number of generations and in years, assuming a mutation rate of 1.2 × 10 ^−8^ per base pair per generation (CONRAD *et al*. 2011; CAMPBELL *et al*. 2012; KONG *et al*. 2012) and a generation time of 24 years (SCALLY and DURBIN 2012) (see Methods). The *T*_*MRCA*_ of the Yoruba population inferred under the four demographic models is of 87 100 generations corresponding to 1.7 million years. The inferred demographic models, with scaling in coalescent units, number of generations and number of years, are shown in Figure 4. The coalescent unit of 67 000 estimated to scale the inferred coalescent times in number of years corresponds to a present effective population size *N*_*e*_(0) of 33 500.

The demography inferred by the stairway plot method for the Yoruba population is a piecewise-constant demography showing much more complex patterns of growth and bottlenecks than the one-parameter models (Figure 2). Moreover, the expected SFS under this inferred demography does not fit well the observed Yoruba SFS (Figure 3B). To understand what could produce such a complex demography, we simulated SFS under a *Linear* growth with the founding time 
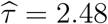
 inferred for the Yoruba population. We simulated three sets of 200 SFS, with respectively 10^3^, 10^4^, and 10^5^ loci, to obtain SFS with more or less noise (solid lines on Figure 5A). We applied the two inference methods to these SFS. The median demographies inferred by the stairway plot method are strongly affected by the noise of the SFS, as shown on Figure 5B. When the number of simulated loci is very large (median of 200 independent demographies inferred with 10^6^ loci), the stairway plot gives a good approximation of the true demography, and the expected SFS under the inferred demography fits the input SFS. However, for smaller numbers of loci (median of 200 independent demographies inferred with 10^5^ loci or less), the stairway plot shows complex patterns of growth and bottlenecks incompatible with the true demography, and the expected SFS under the inferred demographies do not fit the input SFS. On the contrary, the one-parameter method infers a *Linear* demography with a founding time close to the true value for SFS simulated with 10^4^ loci or more (Table 2).

**Table 2:** Inference of the founding time 
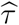
 under the *Linear* model on SFS with noise. Mean, 5% and 95% percentile of the founding time inferred with a *Linear* model. The SFS on which the inference is made are simulated with a founding time *τ* of 2.48, with different number of loci, using the method with topology reconstruction

| Number of loci | 5% percentile | Mean $\hat{\tau}$ | 95% percentile |
| --- | --- | --- | --- |
| $10^3$ | 2.569 | 2.713 | 2.893 |
| $10^4$ | 2.463 | 2.503 | 2.540 |
| $10^5$ | 2.473 | 2.485 | 2.498 |
| $10^6$ | 2.478 | 2.483 | 2.487 |

## DISCUSSION

In this study, we fit the SFS of the Yoruba population with five simple demographic models of growth described by one parameter. Surprisingly, even though these five models are quite distinct in the way they model population growth, fitting them on the Yoruba data results in strongly similar SFS, which all show an excellent goodness of fit with the observed Yoruba SFS. Fitting the same SFS with the stairway plot method (LIU and FU 2015), a model-flexible method which infers a piecewise-constant demography, resulted in a complex demography with several bottlenecks in the last 160 000 years. The poor goodness of fit of the expected SFS under this inferred demography with the Yoruba SFS indicates that this complex demography is not to be trusted and suggests that the way the method estimates the number of change points is too flexible.

The results obtained by the model-constrained and model-flexible methods showed some similarities: the current population size *N*_*e*_(0) of about 30 000 inferred with the stairway plot corresponds roughly to the coalescent unit of 67 000 generations (equivalent to 2*N*_*e*_(0) in the coalescent theory) found with the one-parameter models. Similarly, the *T*_*MRCA*_ of ∼1.7 million years inferred with the one-parameter models seems to match with the last time point of the stairway plot, at about 1.9 million years.

We hypothesize that the complexity of the demography inferred by the stairway plot method is caused by the irregularities of the observed Yoruba SFS. Two concurrent non-exclusive explanations can be put forward for these irregularities. First, they can be due to the sampling and thus be considered as noise that should not be interpreted as evidence for demography. Second, these irregularities could be biologically relevant and result from a very complex demographic history. To assess the impact of noise on the stairway plot method, we tested it on simulated SFS under a *Linear* growth. These SFS were simulated with different numbers of independent loci: the more loci, the less noise in the simulated SFS. The stairway plot inference on these SFS shows that the method is strongly affected by the noise in the SFS simulated data: whereas the demography inferred for a smooth SFS (corresponding to a high number of independent loci) corresponds to the true demography approximated as piecewise constant, the demographies inferred for smaller numbers of loci show complex patterns of bottlenecks and deviate strongly from the true demography. It could be that this method captures the signal contained in these irregularities and infers a demography taking them into account, whereas the one-parameter models fit the global trend of the SFS shape and can thus infer the true demography for much smaller numbers of loci. One solution could be to constrain the number of parameters allowed for model-flexible methods: it seems that determining it by likelihood-ratio test, as it is done in the stairway plot method, is not conservative enough, as it does not prevent from over fitting the noise. If the number of parameters was forced to be small, the method might capture the global trend of the demography and avoid this issue. The SFS reconstructed under the demographies inferred by the stairway plot, however, differ strongly from the input SFS. If the issue was the overfitting of noise, we would expect the reconstructed SFS to t the data more closely. The method is clearly biased by noise on the SFS but it remains unclear why. It would require further investigation to analyze how the different characteristics of this particular method, such as the parametrization of population size history, respond to noise, and what is responsible for this bias.

The five one-parameter demographic models all predict virtually the same SFS for the Yoruba population. Therefore, they also predict the same *T*_*MRCA*_ for the Yoruba population. This *T*_*MRCA*_ of ∼1.3 in coalescent units corresponds, with our scaling of coalescent time based on the number of mutations per site, to ∼1.7 million years. This estimation is similar to results concerning the whole human population, obtained by BLUM and JAKOBSSON (2011) or reviewed in GARRIGAN and HAMMER (2006). Although the commonly admitted date of emergence of the anatomically modern human is around 200 000 years ago, Blum and Jakobsson showed that finding a much older *T*_*MRCA*_ was compatible with the single-origin hypothesis, assuming a certain ancestral effective population size. These ancient times to most recent common ancestor could also be explained by gene flow in a structured ancestral population (GARRIGAN and HAMMER 2006).

Although all five models predict the same *T*_*MRCA*_, the inferred demographies differ substantially between the models (Figure 3A). In the time range further beyond the *T*_*MRCA*_, no information is carried by the sample. Thus, the inferred demographies differ in this time range (Figure 4), making the inferred founding time of the Yoruba population unreliable.

**Figure 4:**
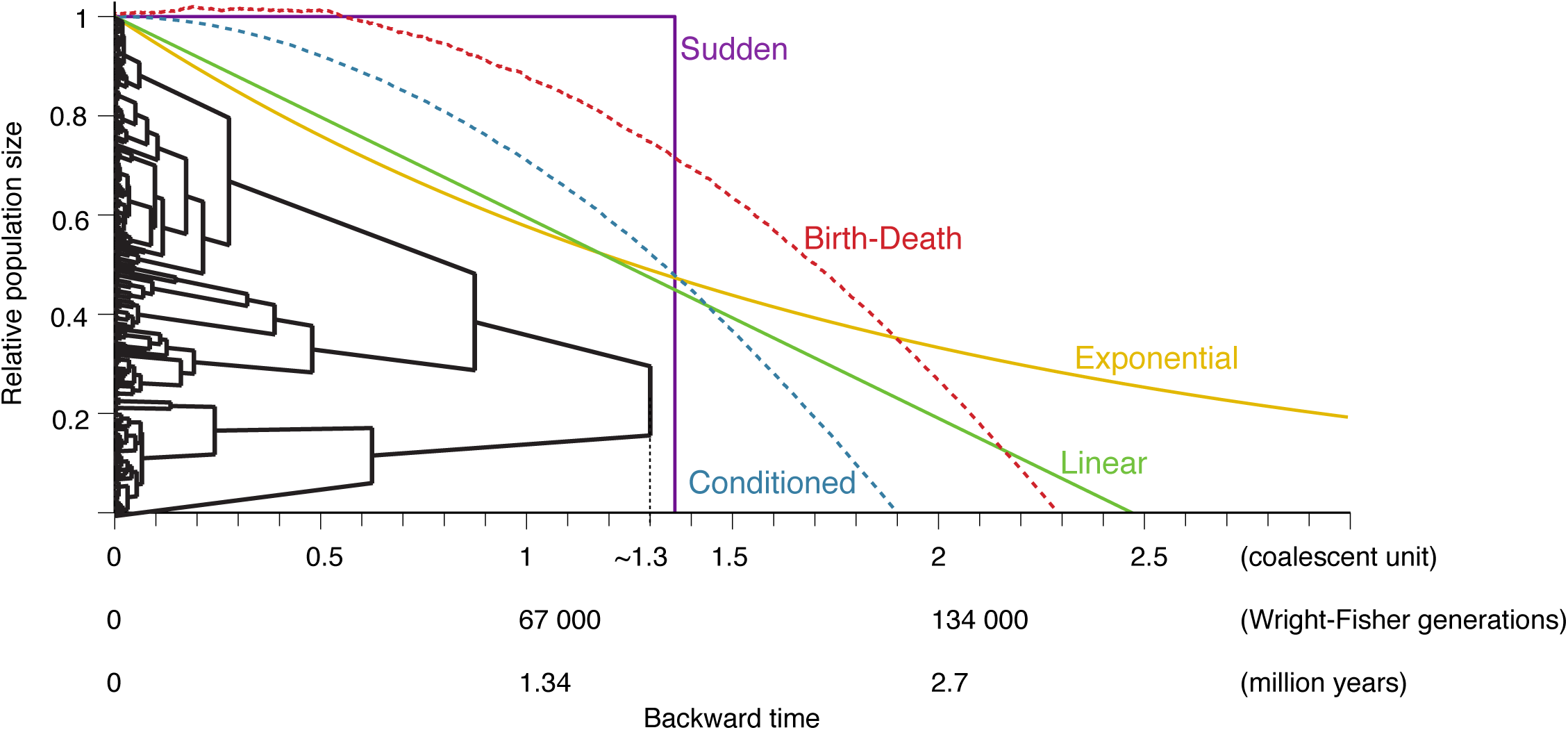
Demographic histories and reconstructed tree estimated from the Yoruba SFS. The tree shown has internode durations *t_k_* during which there are *k* lineages consistent with the SFS (the topology was chosen uniformly among ranked binary trees with 2*n* tips). Time is given in coalescent units, and scaled in number of generations and in millions of years. The demographic histories (solid lines: explicit models, dashed lines: implicit models) are plotted with their optimized 
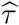
values. See the supplementary methods for details on the demographic histories plotted for the models with implicit demographies (*Birth-Death* and *Conditioned*)

Our results with one-parameter models are reproducible with another model-constrained method, ∂a∂i, which uses different approaches both for the theoretical SFS simulations (diffusion approximation) and the parameter optimization (composite likelihood). This shows that, for models having the same number of parameters, a distance-based approach finds the same ranking of models as a likelihood framework, while being computationally less intensive. Furthermore, the distance-based approach allows for intuitive evidence on the fact that these different models actually all perform very well to fit the Yoruba SFS: the small differences of distance between the best SFS predicted by each model and the observed SFS could be due only to the noise in the observed SFS and thus do not mean that one model is better than another.

Among the five tested demographic models, two pairs of models seem to predict particularly similar SFS (pairs of models with the two smallest values of *d*^2^ in Table 1). First, the *Linear* (L) and *Exponential* (E) growth models predict almost identical SFS for the Yoruba population 
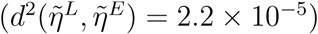

Figure 4 shows that, in the time range where information is conveyed by the mean coalescent tree of the population, *i.e.*, between present time and the *T*_*MRCA*_, these two demographies are very similar. This explains why their SFS are almost indistinguishable, and shows that in this parameter range, it is impossible to distinguish Linear from exponential growth. Second, the SFS predicted under the two models with implicit demography, *Conditioned* (C) and *Birth-Death* (BD), are so similar that they are undistinguishable in Figure 3B

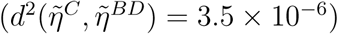
 This raises a question on how these two models, based on different processes — a Wright-Fisher model or a branching process — compare and in particular why their SFS are so similar.

As we compute the distance statistic to optimize the models on normalized SFS, the information of the magnitude of the SFS (often referred to as *θ*, the population mutation rate) is lost. However, as the inferred SFS under the five demographic models all have the same shape, the constant by which they should be multiplied to fit the real, not normalized, Yoruba SFS would be the same for all five models. Thus, this information would not allow to choose which model infers the most realistic value of *θ*.

The outlying first bin of the Yoruba SFS, corresponding to singletons, was removed from our inference because it can be affected by sequencing errors. As the relatively low to moderate coverage of the 1000 Genomes project could also result in an underestimation of doubletons and tripletons, we optimized *τ* masking also these values. It did not change the estimation of 
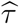
 and thus had no effect on the inferred demographies. As the first bin of the SFS accounts for the mutations that occur in the terminal branches of the coalescent tree, a large part of the excess of singletons can be due to very recent and massive growth. Recent studies with deep sequencing coverage have shown that there is a large abundance of rare variants in human populations (COVENTRY *et al*. 2010; NELSON *et al*. 2012; GAZAVE *et al*. 2014). As the dataset we used for this study had a limited sample size and low coverage, we focused on the inference of demography in the more distant past. Thus, because of both sequencing errors and incompatibility with our one-parameter models, singletons were not taken into account. Our inferences concern the population before this recent and massive growth. It should also be noted that LIU and FU (2015) emphasize that the strength of their method is in capturing recent demographic history. Thus, ignoring singletons, although it is an existing feature of their software, might not be the most appropriate use of the stairway plot.

**Figure 5:**
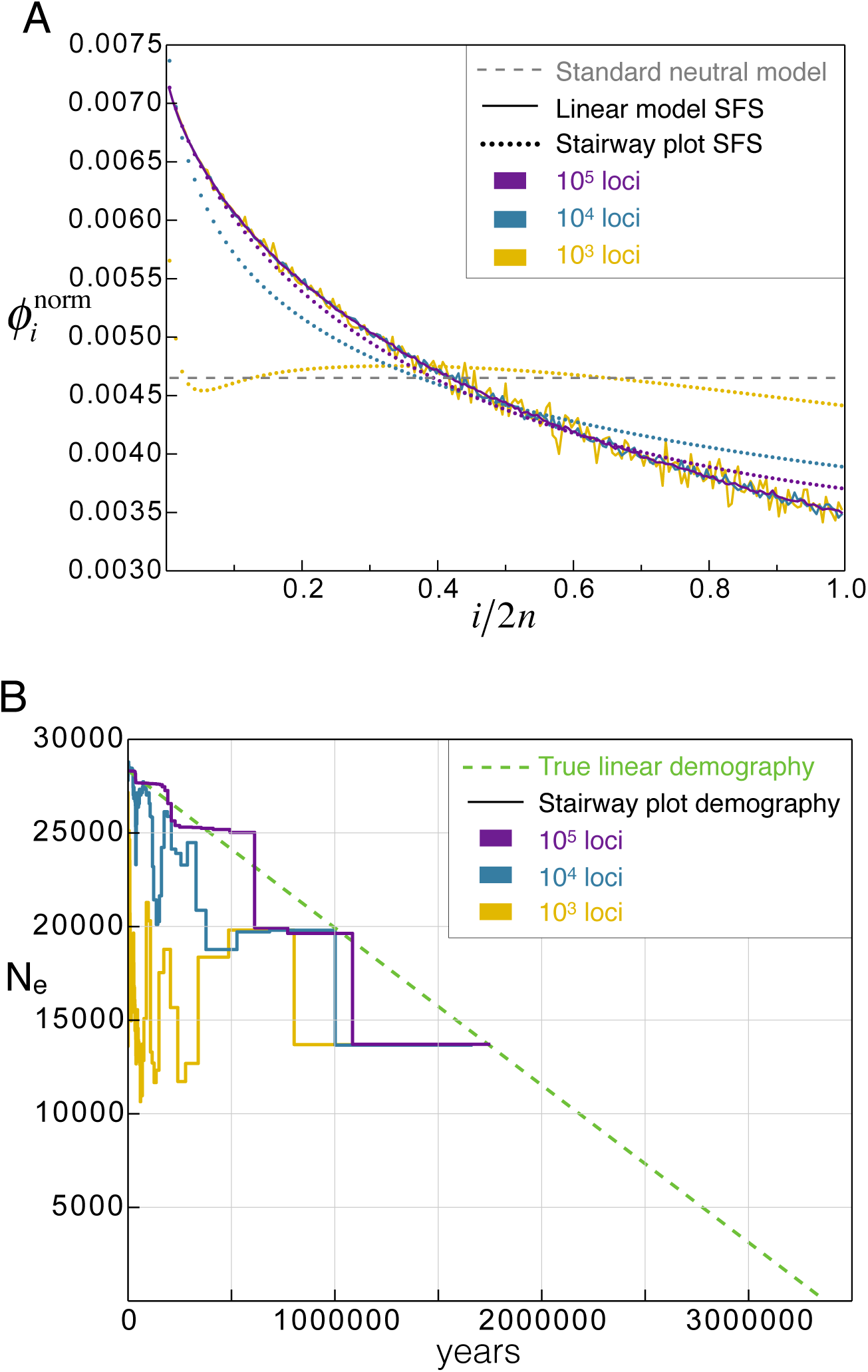
Stairway plot inference of a Linear demography SFS with noise. A) Solid lines: mean of 200 SFS simulated independently under the *Linear* growth model, with either 10^5^ loci (purple), 10^4^ loci (blue) or 10^3^ loci (yellow). Dotted lines: expected SFS under the demography reconstructed by the stairway plot method for different number of loci (same colors than solid lines). The grey dashed line is the expected SFS under the standard neutral model without demography. The SFS are transformed and normalized (see Methods). B) Stairway plot demographic inference: Median of 200 independent demographies inferred with 200 independently simulated SFS for each number of loci (colors match the plot above). The true demography is the green dashed line. The inferred effective size *N_e_* is plotted from present time (0) to the past.

For non-African human population, the SFS based on the 1000 Genomes Project data are not monotonous: their shape is more complex than the SFS of the Yoruba population. Thus, one-parameter models cannot capture the complexity of the demographic histories underlying these types of observed SFS. Even for the Yoruba population, capturing the recent growth event, by taking into account the singletons, would have required adding another parameter. The stairway plot method shows more flexibility and could capture the signal for more complex demographic histories, provided that the number of independent loci is very large so that there is no bias due to noise.

Overall, this study shows that even in the case of a simple demography, the scenario inferred by the stairway plot, a model-flexible method, can show spuriously complex patterns of growth and decline and can predict SFS poorly fitting with the initial SFS data. This might be explained by overfitting of the method to the noise present in the observed SFS, which can be expected for a reasonable number of loci. We also show that simple models described by one parameter can have an excellent goodness of fit to the data and avoid the issue of noise overfitting. The results indicate that the demography of the Yoruba population is compatible with simple one-parameter models of growth, and that the expected *T*_*MRCA*_ of this population can be estimated at ∼1.7 million years. However, the SFS is not sufficient to determine which model better characterizes the Yoruba demographic growth, and estimations of the founding time of the population, that depend on the chosen model, are thus unreliable. More generally, this study illustrates the issue of non-identifiability of demographies based on the SFS of a finite sample.

Our comparison of a model-constrained method using one parameter models with a model-flexible method using a potentially large number of parameters highlights the importance of the model complexity. How many parameters should we use to “properly” characterize a demography? We argue that low complexity models should be tested first. For model-flexible methods, the number of parameters is usually unbounded and determined by successive likelihood ratio tests. This statistical framework implies that a certain risk is taken at each successive step, and that with the repetition of steps, errors can potentially be made. For example, these errors can lead to spurious inferences in noisy data (*i.e*., any real data). We recommend (visually) monitoring the improvement in goodness of fit when adding new parameters on statistical grounds. Examination of the intermediate steps of fitting would likely prevent an unnecessary increase in the model complexity.

## ACKNOWLEDGMENTS

We thank Cécile Delaporte for preliminary work on this project and Simon Boitard, Michael Blum, Konrad Lohse and three anonymous reviewers for useful comments on the manuscript. G.A. and M.L acknowledge support from the grant ANR-12-NSV7-0012 Demochips from the Agence Nationale de la Recherche (France). M.L. is funded by the PhD program ‘Interfaces pour le Vivant’ of UPMC Univ Paris 06. G.A., A.L. and M.L. thank the *Center for Interdisciplinary Research in Biology* for funding.

